# Self-Reported Sleep Relates to Microstructural Hippocampal Decline in β-Amyloid Positive Adults Beyond Genetic Risk

**DOI:** 10.1101/2020.04.28.061184

**Authors:** Håkon Grydeland, Donatas Sederevičius, Yunpeng Wang, David Bartrés-Faz, Lars Bertram, Valerija Dobricic, Sandra Düzel, Klaus P. Ebmeier, Ulman Lindenberger, Lars Nyberg, Sara Pudas, Claire E. Sexton, Cristina Solé-Padullés, Kristine B. Walhovd, Anders M. Fjell

**Affiliations:** Research Group for Lifespan Changes in Brain and Cognition, Department of Psychology, University of Oslo; Department of Radiology and Nuclear Medicine, University of Oslo; Departament de Medicina, Facultat de Medicina i Ciències de la Salut, Universitat de Barcelona, Spain; Lübeck Interdisciplinary Platform for Genome Analytics (LIGA), Institutes of Neurogenetics and Cardiogenetics, University of Lübeck, Lübeck, Germany; Center for Lifespan Psychology, Max Planck Institute for Human Development, Berlin, Germany; Department of Psychiatry, University of Oxford, UK; Max Planck UCL Centre for Computational Psychiatry and Ageing Research, Berlin, Germany, and London, UK; Umeå Center for Functional Brain Imaging, Umeå University, Sweden

**Author notes:** Corresponding author: Håkon Grydeland, Department of Psychology, University of Oslo, PO Box 1094 Blindern, 0317 OSLO, Norway.

**Keywords:** Sleep, hippocampus, β-amyloid, Alzheimer’s disease, memory, mean diffusivity

## Abstract

**Background:** To test the hypothesis that worse self-reported sleep relates to memory decay and reduced hippocampal integrity as indexed by increased intra-hippocampal water diffusion, and that the relations are stronger in the presence of β-amyloid (Aβ) accumulation, a marker of Alzheimer’s disease (AD) pathology.

**Methods:** Two-hundred and forty-three cognitively healthy participants, aged 19-81 years, completed the Pittsburgh Sleep Quality Index, and 2 diffusion tensor imaging sessions, on average 3 years apart, allowing measures of decline in hippocampal microstructural integrity as indexed by increased mean diffusivity. We measured memory decay using delayed recall from the California Verbal Learning Test. ^18^F-Flutemetamol positron emission tomography, in 108 participants above 44 years of age, yielded 23 Aβ positive. Genotyping enabled controlling for *APOE* ε4 status, and polygenic scores for sleep efficiency and AD.

**Results:** Worse global sleep quality and sleep efficiency related to more rapid reduction in hippocampal microstructural integrity over time. Focusing on sleep efficiency, the relation was stronger in presence of Aβ accumulation. Sleep efficiency related to memory decay indirectly via hippocampal integrity decline. The results were not explained by genetic risk for sleep efficiency and AD.

**Conclusions:** Poor self-reported sleep efficiency related to decline in hippocampal integrity, especially in the presence of Aβ accumulation. Poor sleep and hippocampal microstructural decline may partly explain memory decline in older adults with Aβ pathology. The relationships were not explained by genetic risk. Poor self-reported sleep efficiency might constitute a risk factor for AD, although the causal mechanisms driving the of observed associations remain unknown.

## Introduction

Individuals with sleep disturbances have increased risk for Alzheimer’s disease (AD)(1), and accumulation of β-amyloid (Aβ)(2, 3). Aβ is modestly related to memory decline(4), and studies have suggested that relations between Aβ and memory partly depend on sleep(5, 6). A critical role in linking sleep to Aβ and memory may be played by hippocampal integrity. We have previously shown that worse self-reported sleep related modestly to hippocampal atrophy across samples(7). Integrity measured by diffusion tensor imaging (DTI) may detect more subtle microstructural decline(8), and hippocampal mean diffusivity (MD) has demonstrated sensitivity to memory(9). Sleep-hippocampal integrity relationships could also reflect effects of the *APOE* ε4 genotype(10), or of common genetic variation affecting sleep and hippocampus(11). Testing whether worse self-reported sleep relates to memory decline and more rapid reduction of hippocampal integrity while controlling for genetic variation, and whether such relations are stronger in older adults with pathological levels of Aβ, might help us decipher the role of sleep problems in early AD-related pathology.

Here, in 243 cognitively healthy participants aged 19-81 years, we asked whether self-reported sleep characteristics were associated with decline in memory and microstructural (MD) hippocampal integrity over an average of 3 years. We hypothesized worse sleep to be related to stronger decline, particularly in individuals with cortical Aβ accumulation, and also when controlling for *APOE* ε4 and polygenic scores (PGSs) for sleep efficiency and AD(12), respectively. To further assess self-reported sleep relations with memory decline, we also performed a meta-analysis using data from the Lifebrain consortium(13).

## Methods and Materials

### Sample

The sample was drawn from projects consisting of 2-6 study waves at the Center for Lifespan Changes in Brain and Cognition, Department of Psychology, University of Oslo, Norway. The Regional Ethical Committee of Southern Norway approved all procedures, and all participants consented in writing prior to commencement. At baseline, participants were recruited through advertisements. At follow-up, recruitment was by written invitation to the original participants. At both time points, participants underwent health interviews, and were required to be right-handed, fluent Norwegian speakers, and have normal or corrected to normal vision and hearing. Exclusion criteria were history of injury, disease or psychoactive drug use affecting central nervous system function, including clinically significant stroke, serious head injury, untreated hypertension, and diabetes, as well as magnetic resonance imaging (MRI) contraindications. Based the availability of a completed sleep questionnaire, and valid baseline and follow-up anatomical MRI and DTI scans, 251 community-dwelling participants were eligible for inclusion (see **Fig. S1** for an attrition overview). Additional inclusion criteria for the present analyses were (i) valid scores at baseline and follow-up on the long delay free recall of the California Verbal Learning Test (CVLT, see below for details, 7 participants lacked scores at follow-up), and (ii) as in our previous work (14), CVLT long delay free recall change < 60% (one participant excluded). The final sample consisted of 243 participants (62% female, mean baseline age = 54, range: 19-81, see **Table 1** for details).

**Table 1.**
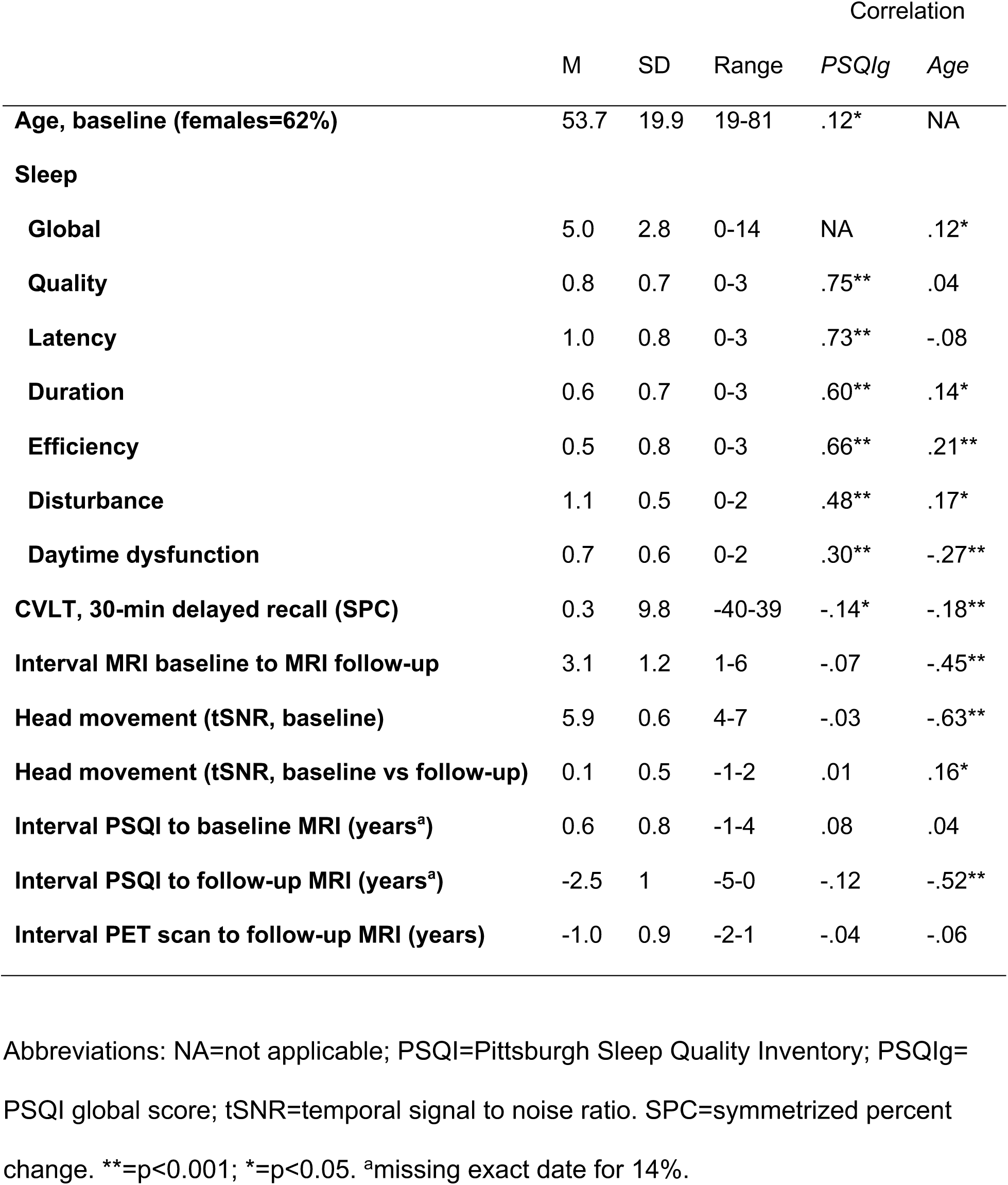
Participants demographics

Participants had full-scale intelligence quotient (IQ) above 85 on the Wechsler Abbreviated Scale of Intelligence (15), except 2 participants aged 64 and 27 years, scoring 79 and 83 at baseline (both scored > 85 on follow-up). On the Mini Mental State Examination (MMS)(16), participants above 40 years of age scored ≥26, except 2 participants aged 80 years scoring 25. All participants who completed the Beck Depression Inventory (BDI) scored ≤16, except 4 participants, aged 24-45 at follow-up, scoring 18-24. Eighty-one participants aged above 68 years completed the Geriatric Depression Scale (GDS)(17), and all scored ≤ 9 except for 7 participants (5 participants aged 71-74 at follow-up, and 2 participants aged 77 and 73 years at baseline which scored at non-depression levels on follow-up). A depression score was missing for 15 participants, either at one time point (13 participants, aged 19-77 years, all scoring ≤ 7 on BDI) or both (2 participants, aged 29 and 58 years). To account for potential influences of particularly depression, we undertook sensitivity analyses (see below). A neuroradiologist evaluated the MRI scans, and all participants were deemed free of significant injuries or pathological conditions.

**Figure 1A** shows the study design. Similar to our previous work on self-reported sleep(14), baseline MRI was administered between 2011 and 2016, and follow-up MRI between 2015 and 2018. PSQI was completed once by each participant, between 2012 and 2017, on average 0.6 (SD=0.8) years after baseline MRI (16 participants completed the PSQI on average 0.4 (SD=0.3) years before baseline MRI, while exact completion date was not available for 34 participants). The memory assessments were performed on average 13 (SD=22) days before the baseline MRI, and 26 (SD=29) days before the follow-up MRI, respectively. Positron emission tomography (PET) scanning was performed once in a subset of participants, between 2015 and 2018, on average 1 (SD=0.9) year before the MRI follow-up.

**Figure 1.**
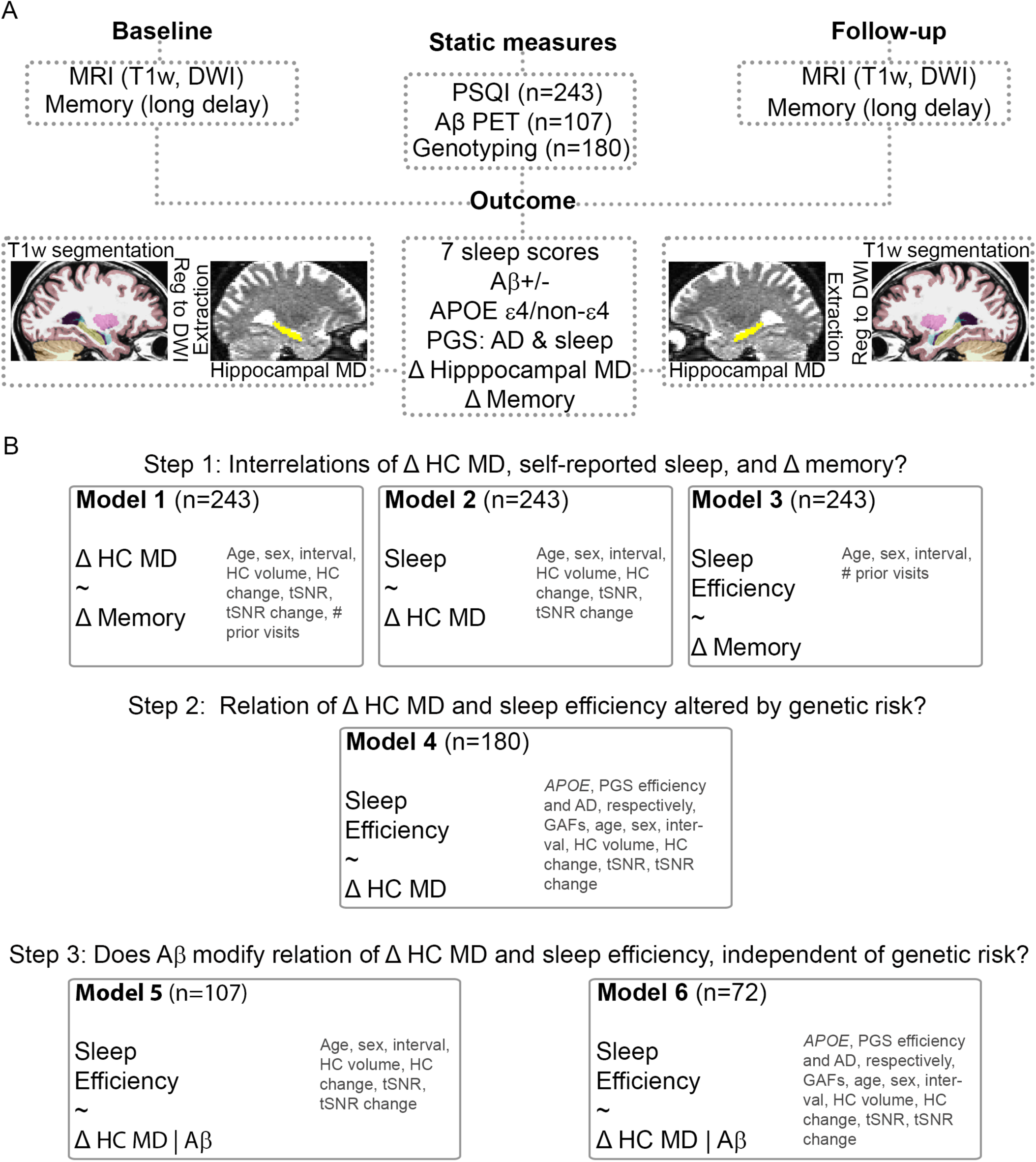
Study overview. A. Study design. B. Main regression models. Covariates are named in dark gray font color. Abbreviations: Age=baseline MRI age; HC=hippocampus; HC volume=baseline hippocampal volume; tSNR=temporal signal to noise ratio, derived from DWI scans (see text for details); PSQI=Pittsburgh Sleep Quality Inventory; Aβ=β-amyloid; PGS=polygenic scores; GAF=genetic ancestry factor; # prior visits=number of prior visits.

### Sleep assessment

To assess sleep, we used the Pittsburgh Sleep Quality Index (PSQI)(18). This self-report index yields one global sleep quality score, which is the sum of the score of 7 components: 1) quality, 2) latency, 3) duration, 4) habitual efficiency, 5) disturbance, 6) use of sleep medication, and 7) daytime dysfunction (see Supplemental Information [SI] for details). We did not evaluate the sixth component as use of medication was an exclusion criterion. Although the PSQI asks about sleep patterns of the last month, here, as in our previous longitudinal work(14), we take the PSQI to reflect relatively stable sleep patterns, an inference for which there is support in adults above 38 years(19, 20).

### MRI acquisition

Diffusion tensor imaging scans were acquired at two Siemens scanners (Siemens Medical Solutions, Erlangen, Germany), a 1.5 T Avanto (n=64, 70% female, mean age (SD, min-max) = 51 (13, 24-77) years), TR/TE=8200/81 ms, FOV=128, 60 diffusion-sensitizing gradients at a b-value of 700 s mm^−2^ and 2 volumes without diffusion weighting (b-value = 0)), and 3T Skyra scanner (n=187, 58% female, mean age (SD, min-max) = 55 (22, 19-81) years), TR/TE=9200/87 ms, FOV=130, 64 diffusion-sensitizing gradients at a b-value of 1000 s mm^−2^ and 1 volume without diffusion weighting. The sequences and scanner were the same across the two time points for each participant.

### Preprocessing

The diffusion-weighted data were analyzed using the FMRIB Software Library (see SI for details), and included susceptibility-induced field correction, and correction for head motion, signal dropout, and eddy current-induced fields(21). After removing nonbrain tissue, and estimating the diffusion ellipsoid properties (the length of the longest, middle, and shortest axes, called eigenvalues), we computed mean diffusivity (MD), defined as the mean of the three eigenvalues.

### Hippocampus segmentation and DTI registration

The T1-weighted image was automatically processed with FreeSurfer software suite (version 6.0.0), independently for each time point, yielding segmentation of left and right hippocampus(22). After co-registering the b=0 volume to the T1-weighted image (see SI for details), inverting the resulting matrix, and applying it to the hippocampal segmentation, we extracted hippocampal MD in diffusion space. To reduce the number of tests, we calculated the average hippocampal MD based on the left and right hippocampus at each time point.

### Memory change

The participants underwent neuropsychological testing including memory assessment via the CVLT. In an effort to minimize practice effects due to repeated testing, we administered alternative versions containing different words and categories at follow-up. We chose the arguably most sensitive measure of hippocampus-dependent memory, namely long delay free recall, that is, the number of correctly recalled words after an approximately 30-minute delay.

### Symmetrized percent change (SPC)

As in our previous longitudinal sleep work(14), we calculated symmetrized percent change (SPC), as symmetrized measures have been shown to be more robust, and with equal or greater statistical power(23). For the average hippocampus value at baseline and follow-up (AH1 and AH2), the SPC was obtained by the following formula: SPC = 100 * (AH2 − AH1)/(AH2 + AH1). The same formula was used to obtain SPC measure for memory change.

### PET acquisition

A total of 108 participants (mean age (SD, min-max)=68.0 (8.7, 44.4-80.8) years) underwent ^18^F-flutemetamol-PET scan, sensitive to Aβ accumulation. Images were acquired on a General Electric Discovery PET/CT 690 scanner at Aleris Hospital and Radiology, Oslo, Norway (see SI for details).

### Genetic data

A subsample of 179 participants (64% females, mean age (SD, min-max) = 53.7 (20.4, 20.1-80.8) years had genome-wide single nucleotide polymorphism (SNP) and manual *APOE* ε4 genotypes available. The PGSs of sleep efficiency and AD were computed (see SI for details) using summary statistics from previously published genome-wide association studies (GWAS) (24, 25). To test for the effect of *APOE* separately from the common genetic variation reflected by the polygenic scores, we estimated *APOE* ε4 counts, coded as 0, 1, or 2 copies of the ε4 allele, and binarized to ε4-non-carrier or ε4-carrier.

### PET pre-processing

We used *PetSurfer*, a set of tools within the FreeSurfer suite, for partial volume correction(26) (see SI for details). Briefly, the procedure yielded PET signal for each of 68 cortical regions. We used cortical regions as Aβ has been reported to appear first in cortex(27). The PET signal in each region was divided by the mean signal of the cerebellum cortex to obtain standardized uptake value ratios (SUVR)(28).

### Aβ status

As common in the literature(28), we dichotomized the SUVR into high or low Aβ groups using a data-driven approach (see SI for details). As previously reported in healthy older participants(28), the optimal model consisted of a 2-distribution model with unequal variance. Participants with a >.5 probability of belonging to the high Aβ distribution were classified as *A*β positive, and the remaining as *A*β negative.

### Meta-analysis of self-reported sleep and memory change

To test the relation between sleep and memory change, we also included data from the Lifebrain consortium(13), yielding a total of 1196 participants. The samples and procedures used are described in detail elsewhere(7), and details of current analysis can be found in SI. The data available in all projects were (i) self-reported sleep scores from one time point, and (ii) memory change score between two time points.

### Statistics

Our main question of a relation between sleep and microstructural hippocampus change was addressed by multiple regression models testing 7 PSQI variables versus hippocampal MD change (see **Figure 1B** for main regression models). To correct for the multiple tests, we adjusted the 7 resulting p-values by applying false discovery rate (FDR) correction (*p*.*adjust* function, R *stats* version 3.6.1). As a proxy measure of head movement during scanning, we calculated temporal signal-to-noise ratio from the diffusion scans(29), which increased with age (R^2^=.40, p<0.001). We included this ratio in all hippocampal MD analyses as covariate of no interest, in addition to age, sex, interval between baseline and follow-up, hippocampus volume at baseline MRI, and difference in movement and hippocampal volume between baseline and follow-up MRI. The latter two covariates were included to (i) assess microstructural effects specifically, and (ii) to correct for volume differences potentially leading to differences in partial volume effects. As participants were drawn from various waves, we included number of prior visits in as a covariate models including memory change to account for potential learning effects. To test whether a relation between sleep and hippocampal MD change was similar across the adult lifespan, we assessed the interaction between the PSQI measure and age. To test for mediation of hippocampal MD change, we performed a mediation analysis across 10000 bootstrapped samples (R package *mediation* v4.5.0)(30). In the Lifebrain consortium data, to test for the relation between sleep and memory change, we calculated partial correlations between sleep and memory change for each sample, correcting for age, sex, and interval between memory tests. We submitted the resulting correlations and corresponding sample sizes to a meta-analysis (R package *meta* v4.9-8). For the analyses including PGSs, the first 3 principal components of the genetic ancestry factors were included as covariates to correct for population substructures. To account for potential influences of depression and cognitive impairment, we undertook two sensitivity analyses. First, we tested whether sleep was related to MD hippocampal change when adding depression, both baseline and change scores, to the covariates in the main analysis (scores from BDI (n=172, median [min-max] baseline age=51 [20-81] years) and GDS (n=43, median [min-max] baseline age=73 [70-81] years) were entered together, with a separate term controlling for depression scale. Second, we excluded excluding the 11 participants with high depression scores, and the 2 participants with a low MMS score, and assessed the similarities with the main results.

## Results

### Sleep and age

Summary measures of the PSQI variables can be found in **Table** 1, together with the correlations between PSQI variables, and between PSQI variables and age. The global score, duration, efficiency, disturbance, and daytime dysfunction, but not quality and latency, showed significant relations with age.

### Microstructural Hippocampal Change and Memory change

Hippocampal microstructural change related to memory change (p<0.001, R^2^=.041, **Fig. 2A**) after accounting for covariates. Higher hippocampal MD change values, interpreted as reduced structural integrity(31), related to more memory decline. Results were similar across the age range (memory change and age interaction p=0.661), and when adding baseline IQ to the covariates (memory change p<0.001).

**Figure 2.**
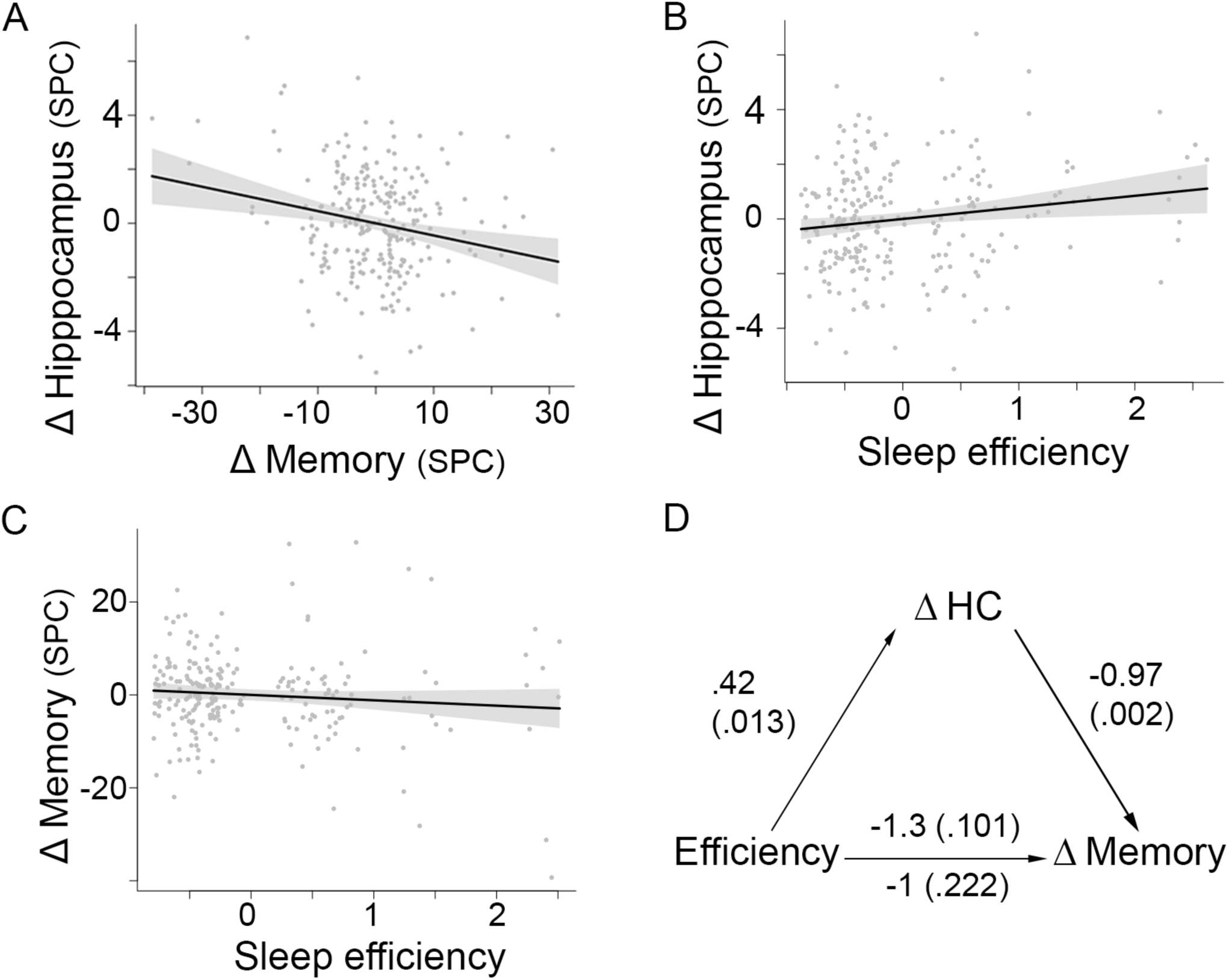
Sleep, and Decline in Microstructural Hippocampal, and Memory. A. Decline in memory related to MD increase in hippocampus (decline in structural integrity). Values are residuals after regressing out covariates (see also **Fig. 1B**). B. Sleep efficiency related to hippocampal MD change, independently of hippocampal volume and hippocampal volume change, after FDR-correction for multiple comparisons. C. Sleep efficiency correlated weakly with memory change (partial r=-.11, correcting for age at baseline, interval, sex, and number of prior visits). D. Average causal mediation effect, that is, the indirect effect of sleep on memory via hippocampus, was −0.41 (p=0.010).

### Sleep and Microstructural Hippocampal Change

We found a relation between hippocampal MD change and (i) the global PSQI score (FDR-corrected p (p_FDR_)<0.05, R^2^=.024, **Fig. S2**), uncorrected p (p_uncorr_)=0.012), and (ii) sleep efficiency (p_FDR_<0.05, R^2^=.023, p_uncor_r=0.013, **Fig. 2B**). The relations revealed that participants with poorer sleep (higher scores) showed more increase in hippocampal MD, independently of hippocampal volume and hippocampal volume change. The relations did not differ across the age range (p_uncorr_ ≥0.319). As the sleep efficiency measure conveys more specific information regarding sleep than the global sleep score, we selected this measure for further analyses.

### Sleep Efficiency and Memory change

Poor sleep efficiency was not strongly related to memory decline (p=0.100, R^2^=0.010, partial r=-.11, **Fig. 2C**). To test if this result accurately reflected the true relation, we performed a meta-analysis in 5 samples from the Lifebrain consortium. This analysis yielded a correlation of −.08 (95% confidence intervals (CI) [-.13, −.02]), Z=-2.70, p=0.007). The partial correlation obtained in the main sample was within this confidence interval, suggesting a modest relation between sleep efficiency and memory change may exist, but needing a larger sample to detect it.

### Sleep Efficiency, Microstructural Hippocampal Change, and Memory Change

To test for hippocampal MD change as a mediator, we ran a mediation analysis (**Fig. 2D**). The unstandardized indirect effect on memory change from sleep efficiency via hippocampal MD change was 0.42 × −0.94 = −0.41, similar to the median bootstrapped unstandardized indirect effect of −0.41 (p=0.010, 95% CI [-0.90, −0.06], ρ at which the effect equals 0 was −0.19). The median direct effect estimate, from sleep efficiency to memory change controlling hippocampal MD change, was −1 (p=0.222). These results suggested that hippocampal MD change partly mediated the relation between sleep efficiency and memory change.

### Sleep Efficiency, Hippocampal Change, and Genetic Effects

The sleep efficiency PGS did not relate to worse self-reported efficiency (**Fig. S3A**, partial r=-.04, p=0.619). Lower genetic propensity for efficient sleep related more strongly, but still very modestly, to hippocampal MD change (**Fig. S4A**, partial r=-.13, p=0.087). For *APOE*, a total of 70 participants carried one or two ε4 alleles. *APOE* ε4 status was not related to sleep efficiency (r=-.05, p=0.526), or hippocampal MD change (r=.08, p=0.307). Re-running the main analysis above adding the sleep efficiency PGS and *APOE* ε4 status, PSQI sleep efficiency still related to hippocampal MD change (p=0.031).

Higher genetic risk for AD was not related to worse sleep efficiency, that is, higher PSQI scores (**Fig. S3B**, partial r=.03, p=0.739), or lower hippocampal MD change (**Fig. S4B**, partial r=-.06, p=0.432). Re-running the main model adding the AD PGS and *APOE* ε4 status, sleep efficiency still related to hippocampal MD change (p=0.023).

### Sleep Efficiency, Hippocampal Change, and Aβ

Of 108 participants with PET data, 23 participants were classified as Aβ positive (**Fig. S5)**. We found a stronger relation between sleep efficiency and hippocampal MD change in participants classified as Aβ positive (efficiency × Aβ interaction term p=0.022, **Fig. 3A**). Including baseline IQ yielded a similar result (interaction term p=0.017). The Aβ positive participants did not show different sleep efficiency (p=0.913), hippocampal MD decline (p=0.932), or memory decline (p=0.680). When repeating the analysis in the Aβ positive and negative groups separately, we observed a relation between sleep efficiency and hippocampal MD change only in the Aβ positive (p=0.019), but not in the Aβ negative subgroup (n=85, p=0.361).

**Figure 3.**
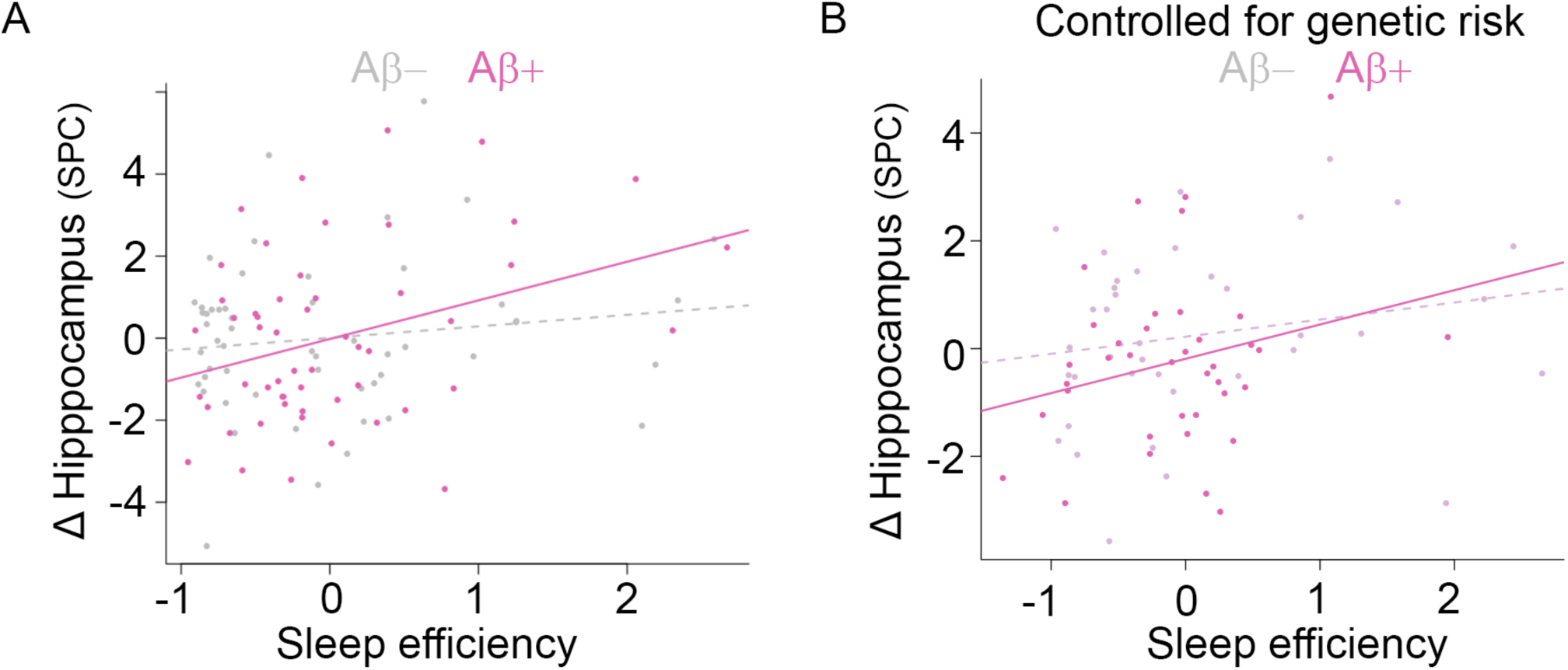
Sleep efficiency, microstructural hippocampal decline, and Aβ accumulation. A. Efficiency related more strongly to microstructural hippocampal decline in participants with signs of cortical Aβ accumulation. B. This relation remained when controlling for APOE ε4 status and PGS for sleep efficiency and AD, respectively.

### Sleep Efficiency, Hippocampal Change, Aβ, and Genetic Effects

A subsample of 76 participants (mean (SD) age=69.3 (8.2) years, min-max 44-81 years) had both Aβ and genotype data, and 24 of these participants had one or two *APOE* ε4 alleles. We included the *APOE* ε4 status and the PGS for sleep efficiency and AD, respectively, to the initial model. The results demonstrated (i) that sleep efficiency still related to hippocampal MD change differently for Aβ negative and positive participants (p=0.015, **Fig. 3B**), and (ii) an effect of the PGS for sleep efficiency on hippocampal MD change (p=0.028), with higher propensity of efficient sleep showing less MD hippocampal decline. The AD PGS showed a very weak effect (p=0.087), while the *APOE* genotype showed no effect (p=0.537).

### Sensitivity analyses

We verified that (i) when controlling for levels of depression (both baseline and change scores), sleep efficiency still related to MD hippocampal change (sleep efficiency p=0.004), and (ii) when excluding the participants with depression and MMS scores beyond threshold values, the results remained highly similar. That is, sleep efficiency related to hippocampal MD change (p=0.011), and weakly to memory change (partial r = −.09), while the relation between sleep efficiency and hippocampal MD change differed depending on cortical Aβ accumulation (p=0.012).

## Discussion

The results indicate that sleep efficiency and hippocampal microstructural decline are related, particularly in presence of cortical Aβ accumulation. This relation does not appear to be explained by *APOE* genotype, or polygenic scores for sleep efficiency or AD. Sleep efficiency also related to memory reduction indirectly via hippocampal integrity decline. Although we cannot rule out that these Aβ-related correlations stem from unexplored factors such as tau deposition in the medial temporal lobes, one possibility is that Aβ accumulation constitutes a vulnerability when present alongside reduced hippocampal integrity, leading to lower sleep efficiency and decline in episodic memory.

As the observed hippocampal effects were independent of baseline hippocampal volume and volume change, microstructural change in the hippocampus might be a particularly sensitive marker of early decline, complementary to atrophy. In support of this hypothesis, two previous studies of 147 (overlapping with the current sample)(14) and 66(32) participants, respectively, did not observe relations between sleep and hippocampal volume or atrophy, while one cross-sectional study including 1201 young adults reported associations between right hippocampal MD and sleep quality (but not sleep duration). These findings also suggest larger samples is needed to detect the sleep-atrophy relations. In support of this notion, in 3105 cognitively normal participants aged 18-90 years, including participants from the present sample, we found that poorer sleep efficiency, as well as sleep quality, problems, and daytime tiredness, were related to greater hippocampal volume loss(7). The current finding supports this relation between self-reported sleep and hippocampal change but extends previous knowledge by revealing independent intra-hippocampal reductions in microstructural integrity.

The mechanisms of microstructural hippocampal decline remain unclear, but may relate to decay of the dendritic architecture. In mice, hippocampal dendritic spine densities have been shown to be reduced both in aging(33), and after sleep deprivation(34). Hippocampal dendritic decay might also underlie the relations observed here between microstructural hippocampus decline and memory reductions. Hippocampal dendritic spine loss in mice has been related to memory defects(35). Over time, loss of spines and synapses might promote larger dendritic disruptions, which in mice have been detected via intra-hippocampal DTI, and linked with memory impairments(36). In humans, these speculations could be tested using ultrahigh-resolution DTI(37).

The relation between sleep and Aβ appears reciprocal, as Aβ both increases after sleep deprivation(38), and increases wakefulness and alters sleep patterns(39). Here, although Aβ status did not relate to sleep efficiency (in contrast to (2, 40)) or hippocampus decline alone (as in (41), but see (42)), the sleep-hippocampal decline relation was stronger in the Aβ positive. Echoing these result, in a separate sample of older adults, we recently observed that tau and YKL-40, a biomarker of inflammation and astroglial activation, related more strongly to the PSQI global score in Aβ positive than in Aβ negative (43). These results raise the possibility that Aβ accumulation co-occurring with other adverse signs such as hippocampal decline or inflammation, signals sleep problems not observed with Aβ accumulation alone.

As we observed that the effect of sleep efficiency on memory decline was mediated by higher hippocampal diffusivity, we hypothesize that hippocampal decline, when concomitant with Aβ accumulation, causes sleep problems, here in the form of poorer sleep efficiency. A potential mechanism might be that the hippocampal decline causes altered brain oscillations affecting sleep(44). The current data does not allow inferences that rule out the reverse causality, of sleep affecting hippocampal decline. We find this reverse pattern less likely as effects were specific for sleep efficiency, rather than sleep duration or quality, both more likely drivers of possible sleep-generated causal effects. We cannot rule out that a variable not assessed here can account for the observed correlations(30). For instance, sleep spindles have been linked to both Aβ and tau(3), and tau potentially related to AD is seen first in the locus coeruleus(45). Activity in this region can alter sleep spindles, affecting memory consolidation(46). To tease out potential causal pathways, studies could include Aβ-negative participants with healthy sleep patterns and no signs of hippocampal decline, and follow them to assess changes in sleep patterns, hippocampal integrity, Aβ status, memory performance, tau, and measures of neuroinflammation such as YKL-40 or sTREM2. Intervention studies targeting for instance hippocampal-dependent cognition(47), and investigating similar markers could be a less costly strategy.

The relations remained after controlling for genetics risk indexed by PGSs for sleep efficiency and AD, respectively, as well as the presence of the *APOE* ε4 allele. *APOE* ε4 is a risk factor for AD(48), and healthy ε4 carriers have shown more pronounced longitudinal hippocampal atrophy(49) and worse sleep quality(50). In contrast, there are reports of no relation between hippocampal atrophy in cognitively healthy adults and AD genes from an exploratory GWAS(51), and of lack of *APOE* effects on hippocampal volume(52). Both for the *APOE* genotype and the PGSs, we observed weak relations with sleep efficiency and hippocampal change, respectively. These low correspondences must be resolved in larger samples before we can draw the conclusion that the relation between sleep efficiency and hippocampal decline is partly independent of genetics.

Limitations of this study include the use of a self-report measure of sleep, at one time point, instead of objective measures such as activity monitors, or polysomnography, collected repeatedly. Although self-reported sleep measures might provide more representative data on sleep than a single-night polysomnography(53), a relatively modest correlation of 0.47 has been reported between reported and measured sleep duration(54). In future studies, a likely key is repeated measurements of sleep patterns, assessment of potential underlying sleep disorders, and the inclusion of other biomarkers. Inclusion of such markers would also improve analyses of mediation, which here does not establish causality. As the sleep efficiency PGSs stem from a GWAS using activity monitors(25), the inclusion of objective sleep measures could shed further light on the relative contribution of sleep genetics and sleep behavior. Although the current sample is relatively large, the potentially complex interplay between Aβ positivity and other markers of relatively low prevalence highlights the need for even larger sample sizes.

The results indicate that hippocampal microstructural decline related to sleep efficiency, and episodic memory change in cognitively normal older adults, particularly in Aβ positive participants. This relation was not readily explained by genetic effects. Poor self-reported sleep efficiency might constitute a risk factor for AD, and future studies need to address why sleep is related to more hippocampal decline in Aβ positive older adults even without dementia.

## Acknowledgements

The Lifebrain project is funded by the EU Horizon 2020 Grant: ‘Healthy minds 0–100 years: Optimizing the use of European brain imaging cohorts (“Lifebrain”)’. Grant agreement number: 732592.

LCBC: The European Research Council under grant agreements 283634, 725025 and 313440, as well as the Norwegian Research Council and The National Association for Public Health’s dementia research program, Norway. Betula: a scholar grant from the Knut and Alice Wallenberg (KAW) foundation to L.N. Barcelona: Partially supported by a Spanish Ministry of Science, Innovation and Universities (MICIUN) grant to D-BF [grant number RTI2018-095181-B-C21 (AEI/FEDER, UE)], and an ICREA Academia 2019 Award; by the Walnuts and Healthy Aging study (http://www.clinicaltrials.gov; Grant NCT01634841) funded by the California Walnut Commission, Sacramento, California. BASE-II has been supported by the German Federal Ministry of Education and Research under grant numbers 16SV5537/16SV5837/16SV5538/16SV5536K/01UW0808/01UW0706/01GL1716A/01 GL1716B, and from the European Research Council under grant agreement 677804. Work on the Whitehall II Imaging Substudy was mainly funded by Lifelong Health and Well-being Programme Grant G1001354 from the UK Medical Research Council (“Predicting MRI Abnormalities with Longitudinal Data of the Whitehall II Substudy”).

This article was published as a preprint on bioRxiv: https://doi.org/10.1101/2020.04.28.061184

