## Supplemental Information for "Self-Reported Sleep Relates to Microstructural Hippocampal Decline in β-Amyloid Positive Adults Beyond Genetic Risk"

Max Planck Institute for Molecular Genetics, Germany (L.B.)

### Supplementary Methods

#### Sample

The sample was drawn from projects consisting of 2-6 study waves at the Center for Lifespan Changes in Brain and Cognition, Department of Psychology, University of Oslo, Norway. The Regional Ethical Committee of Southern Norway approved all procedures, and all participants consented in writing prior to commencement. At baseline, participants were recruited through advertisements. At follow-up, recruitment was by written invitation to the original participants. At both time points, participants underwent health interviews, and required to be right-handed, fluent Norwegian speakers, and have normal or corrected to normal vision and hearing. Exclusion criteria were history of injury, disease or psychoactive drug use affecting central nervous system function, including clinically significant stroke, serious head injury, untreated hypertension, and diabetes, as well as magnetic resonance imaging (MRI) contraindications. Based the availability of a completed PSQI, and valid baseline and follow-up anatomical MRI and DTI scans, 251 community-dwelling participants were eligible for inclusion (see **Fig. S1** for attrition of participants). Additional criteria for being included in the present analyses were 1) valid scores on the long delay free recall of the California Verbal Learning Test (CVLT, see below for details) at baseline and follow-up (7 participants lacked data at follow-up), and, as in our previous work (14), CVLT long delay free recall change < 60% (one participant was excluded due to this criterion).

### Sleep assessment

To assess sleep, we used the Pittsburgh Sleep Quality Index (PSQI)(18). This self-report index yields one global sleep quality score, which is the sum of the score of 7 components: 1) quality, 2) latency, 3) duration, 4) habitual efficiency, 5) disturbance, 6) use of sleep medication, and 7) daytime dysfunction. In PSQI, efficiency is calculated as sleep duration (hours slept) divided by the number of hours spent in bed, times 100, and then given score of 0-3 for >85%, 75-84%, 65-74%, and <65%, respectively(18). We did not evaluate the sixth component as use of medication was an exclusion criterion. Although the PSQI asks about sleep patterns of the last month, here, as in our previous longitudinal work(14), we take the

PSQI to reflect relatively stable sleep patterns, an inference for which there is support in adults above 38 years(19, 20). In line with this premise, the PSQI self-report was not necessarily completed in close proximity to the baseline MRI scan.

### MRI acquisition

Diffusion tensor imaging scans were acquired at two Siemens scanners (Siemens Medical Solutions, Erlangen, Germany), a 1.5 T Avanto (n=64, 70% female, mean age (SD, min-max) = 51 (13, 24-77) years), TR/TE=8200/81 ms, FOV=128, 60 diffusion-sensitizing gradients at a b-value of 700 s $mm^{-2}$ and 2 volumes without diffusion weighting (b-value = 0)), and 3T Skyra scanner (n=187, 58% female, mean age (SD, min-max) = 55 (22, 19-81) years), TR/TE=9200/87 ms, FOV=130, 64 diffusion-sensitizing gradients at a b-value of 1000 s $mm^{-2}$ and 1 volume without diffusion weighting. The sequences and scanner were the same across the two time points for each participant.

### Preprocessing

The diffusion-weighted data were analyzed using the FMRIB Software Library, and included susceptibility-induced field correction with *topup* (Andersson et al., 2003), and correction for head motion, signal dropout, and eddy current-induced fields using *eddy* [23, 24]. After removing nonbrain tissue, eigenvalue maps were computed. We defined mean diffusivity (MD) as the mean of the three eigenvalues. We employed a DTI-derived measure as results indicate that DTI can detect subtle effects in cellular microstructure, which has previously proven sensitive to age-related lifespan changes[25], and particularly hippocampal MD has been shown sensitive to memory[10, 11].

### Hippocampus segmentation and DTI registration

The T1-weighted image was automatically processed with FreeSurfer software suite (version 6.0.0), independently for each time point (as no co-registration across time points was necessary), yielding segmentation of left and right hippocampus[26]. To extract MD from the hippocampi in native DTI space for each participant, a B=0 volume from the diffusion data was registered to the T1-weighted image in FreeSurfer space with a within-subject, cross-modal registration using a boundary-based cost function constrained to be 6 degrees of freedom[27]. The resulting registration matrix was inverted, and applied to the segmentation of the left and right hippocampus, yielding hippocampus masks in native diffusion space. The masks were binarized using *mri_binarize* at a minimum voxel threshold of 1, for the most restricted masks compared with lower thresholds. To reduce the number of tests, we calculated the average hippocampal MD based on the left and right hippocampus at each time point.

### Memory change

The participants underwent neuropsychological testing including memory assessment via the California Verbal Learning Test, second edition (CVLT-II)[28]. In an effort to minimize practice effects due to repeated testing, we administered alternative versions containing different words and categories at follow-up. From this word learning test, we chose the arguably most sensitive measure of hippocampus-dependent memory[29], namely long delay free recall, that is, the number of correctly recalled words after an approximately 30-minute delay (during which other cognitive tests were performed).

### Symmetrized percent change (SPC)

As in our previous longitudinal sleep work[19], we calculated symmetrized percent change (SPC), as symmetrized measures have been shown to be more robust, and with equal or greater statistical power[30]. For the average hippocampus value at baseline and follow-up (AH1 and AH2), the SPC was obtained by the following formula: SPC = 100 * (AH2 - AH1)/(AH2 + AH1). The same formula was used to obtain SPC measure for hippocampal volume and memory change.

### PET acquisition

A total of 107 participants (mean age (SD, min-max)=68.0 (8.7, 44.4-80.8) years) underwent $^{18}$F-flutemetamol-PET scan, sensitive to Aβ accumulation[31]. Images were acquired on a General Electric Discovery PET/CT 690 scanner at Aleris Hospital and Radiology, Oslo, Norway. A low-dose computerized tomography scan was first performed for subsequent attenuation correction of the PET scan. Participants were injected with 200±20 MBq $^{18}$F-flutemetamol as a bolus and examined 90 minutes later. Three-dimensional dynamic data were acquired in list mode for 20 minutes, with the following parameters: 47 image planes, voxel size = 1.33 mm x 1.33 mm x 3.27 mm, field of view = 256 mm. The images were reconstructed using the VUEPoint HD Sharp iterative reconstruction algorithm. This algorithm adds resolution recovery in an iterative reconstruction loop by incorporating information about the PET detector response which improves resolution and contrast recovery compared with traditional analytic methods[32]. We used 4 iterations, 16 subsets, time of flight, and a full width at half maximum Gaussian post-filter of 3 mm. As we were interested in the gross tracer uptake, we binned the data into a single frame, and submitted this static PET image to further pre-processing and value extraction.

### Genetic data

A subsample of 179 participants (64% females, mean age (SD, min-max) = 53.7 (20.4, 20.1-80.8) years had genome-wide single nucleotide polymorphism (SNP) and manual *APOE* ε4 genotypes available. Buccal swab and saliva samples were collected for DNA extraction followed by genome-wide genotyping using the “Global Screening Array” (Illumina, Inc.). A*POE* ε4 (rs429358) status was determined using TaqMan (Thermo Fisher Scientific, Inc.) chemistry. Detailed information on DNA collection, quality control, genotyping, and imputation has been reported elsewhere[33]. The PGSs of sleep efficiency and AD were computed using summary statistics from previously published genome-wide association studies (GWAS) [34, 35]. These statistics were based on SNPs with p-values <0.01 in the respective GWAS, except for variants located in the extended MHC region (build hg19; chr6:25,652,429-33,368,333), where we included the most significant SNP. After removing the *APOE* gene region (build hg19; chr19:44,909,011-45,462,650) for which we used the manually derived ε4 (rs429358) genotypes instead, we used the software PLINK[36] to implement the following steps: (i) clumping of the GWAS summary statistics by the –clump option with parameters --clump-p1 1.0 –clump-p2 1.0 –clump-kb 500 –clump-r2 0.1. The linkage disequilibrium (LD) structure was based on the European subpopulation from the 1000 Genomes Project Phase3[37]. (ii) Deriving PGSs for our sample using the –score function. To control for population substructures, we computed the genetic ancestry factors using principal components methods[38], and included only participants of European ancestry in the genetic subsample analysis. The PGS for sleep efficiency was based on a genome-wide association study using accelerometer-derived mean sleep efficiency (calculated as proportion of sleep period time-window classified as sleep)[35], and in our sample a higher PGS reflected a higher genetic propensity towards more efficient sleep. The AD PGS was based on a genome-wide meta-analysis of clinically diagnosed AD and AD-by-proxy (based on parental diagnoses)[34], and in our sample a higher PGS reflected a higher AD risk. To test for the effect of *APOE* separately from the common genetic variation reflected by the polygenic scores, we estimated *APOE* ε4 counts by determining the haplotypes of the two SNPs rs7412 and rs429358[39, 40], coded as 0, 1, or 2 copies of the ε4 allele, and binarized to ε4-non-carrier or ε4-carrier.

### PET pre-processing

We used *PetSurfer*, a set of tools within the FreeSurfer suite, for partial volume correction. Specifically, for each participant, we registered the static PET image to the anatomical T1-weighted image using boundary-based registration[27]. This registration was inverted to get a high-resolution segmentation (upsample factor = 2) from the high-resolution MRI space in PET space, and simultaneously perform the partial volume correction with the Symmetric Geometric Transfer Matrix method, as recommended when using regions of interests (instead of vertex-wise) approach[41, 42]. This procedure yielded PET signal for each of the 68 cortical regions in Desikan-Killiany atlas[43]. We used cortical regions as Aβ has been reported to appear first in cortex[44]. The PET signal in each cortical region was divided by the mean signal of the cerebellum cortex to obtain standardized uptake value ratios (SUVR)[45].

### Aβ status

As common in the literature[45], we dichotomized the SUVR into high or low Aβ groups using a data-driven approach. We ran a principal component analysis on SUVR from the 68 cortical regions using the *prcomp* function (R package *stats* v3.6.1, values were zero-centered and scaled to have unit variance), and extracted the first component (which explained 66.7% of the variance, while, for comparison, the second component explained 7%). The cut-off between groups was determined using Gaussian mixture modeling (R package *mclust* v5.2). We fitted 18 models, ranging from 1 to 9 mixtures, allowing for either equal or unequal variance, and selected the model with the lowest Bayesian information criterion value. As previously reported in healthy older participants[45], the optimal model consisted of a 2-distribution model with unequal variance. Participants with a >.5 probability of belonging to the high Aβ distribution were classified as *Aβ positive*, and the remaining as *Aβ negative*.

### Meta-analysis of self-reported sleep and memory change

To test the relation between the relevant sleep variable (see below for selection criteria) and memory change, we also included data from the Lifebrain consortium (http://www.lifebrain.uio.no/)[15], an EU-funded (H2020) project including participants from several major European brain studies: Berlin Study of Aging-II (BASE-II)[46, 47], the BETULA project[48], University of Barcelona brain studies[49-51], and Whitehall-II[52], yielding a total of 1196 participants. The samples and procedures used are described in detail elsewhere[8]. The data available in all projects were (i) self-reported sleep scores from one time point, and (ii) memory change score between two time points. All subsamples used the PSQI for sleep evaluation, except the Betula sample, which used the Karolinska Sleep Inventory (for details of conversion to PSQI scores, see[8]). The following memory tests were used: 30-minutes delayed free recall from the Verbal Learning and Memory Test (BASE-II), an immediate free recall of sentences (Betula), 30-minutes delayed recall from the Rey Auditory Verbal Learning Test (Barcelona), a short-term 20 word free recall test (Whitehall-II)[53].

### Statistics

Our main question of a relation between sleep and microstructural hippocampus change was addressed by multiple regression models testing each of the 7 PSQI variables (1 global and 6 components) versus hippocampal MD change. To correct for the multiple tests in this analysis, we adjusted the 7 resulting p-values by applying false discovery rate (FDR)[54] correction and the *p.adjust* function (R *stats* version 3.6.1). Head movement is a potential important confound in brain imaging studies. As a proxy measure of head movement during MRI diffusion scanning, we calculated temporal signal-to-noise ratio (tSNR) from the scans[55]. As expected, head movement increased with age ($R^2$=0.40, p<0.001), and we included tSNR in all hippocampal analyses to account for movement-related artifacts (see **Figure 1B** in the main text for overview of main regression models and corresponding covariates). We included interval between baseline and follow-up as covariate of no interest, in addition to age, and sex. As participants were drawn from various waves, we included number of prior visits as a covariate to account for potential learning effects on the memory task (please note that, as mentioned above, different versions were used at each visit). In the analyses of hippocampal MD change, we also included as covariates as no interest hippocampus volume at baseline MRI, and difference in movement and hippocampal volume between baseline and follow-up MRI. These covariates were included to (i) assess microstructural effects specifically, and (ii) to correct for volume differences potentially leading to differences in partial volume effects. For the one hippocampal volume analysis run to compare with previous studies, we also included estimated intracranial volume[56]. To test whether the relations between sleep and hippocampal MD change was similar across the adult lifespan, we assessed the interaction between the PSQI measure and age. To test for mediation of hippocampal MD change, we performed a mediation analysis across 10000 bootstrapped samples (R package *mediation* v4.5.0)[57, 58]. To test for the relation between sleep and memory change in the Lifebrain consortium data, partial correlations between sleep and memory change were calculated for each sample, correcting for age, sex, and interval between memory tests. We submitted the resulting correlations and corresponding sample sizes to a meta-analysis (R package *meta* v4.9-8). To illustrate the individual data points, and to provide a general measure of effect size, we extracted hippocampal MD SPC values and the PSQI measure of interest, removed the effects of the nuisance regressors, and plotted the resulting residuals, and reported their $R^2$. For the analyses including PGSs, the first 3 principal components of the genetic ancestry factors were included as covariates to correct for population substructures. To account for potential influences of depression and cognitive impairment, we undertook two sensitivity analyses. First, we tested whether sleep was related to MD hippocampal change when adding depression, both baseline and change scores, to the covariates in the main analysis (scores from BDI (n=172, median [min-max] baseline age=51 [20-81] years) and GDS (n=43, median [min-max] baseline age=73 [70-81] years) were entered together, with a separate term controlling for depression scale. Second, we excluded excluding the 11 participants with high depression scores, and the 2 participants with a low MMS score, and assessed the similarities with the main results.

### Supplementary Figures

**Figure S1**. Attrition.

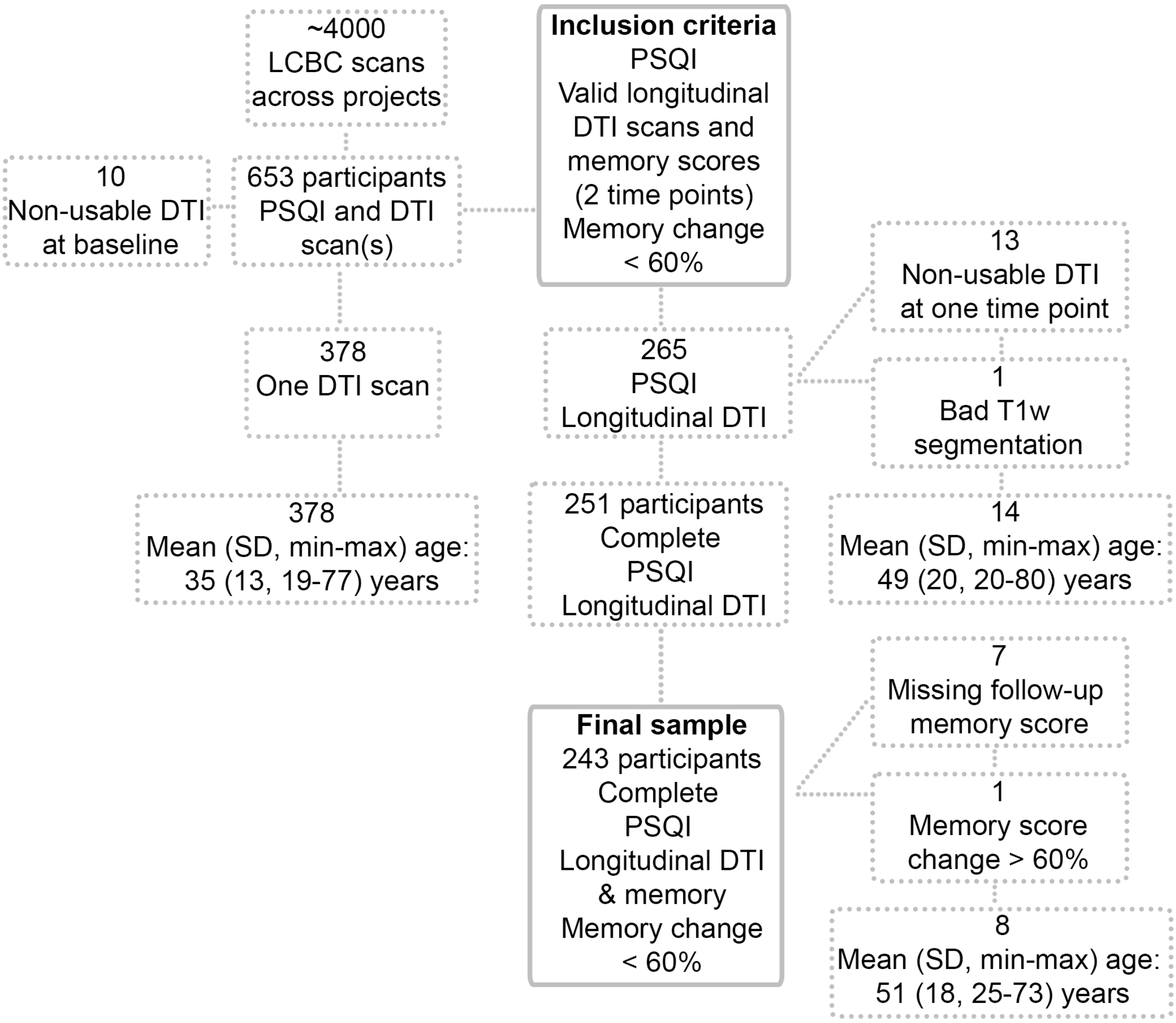

**Figure S2**. Global sleep quality related to hippocampal MD change.

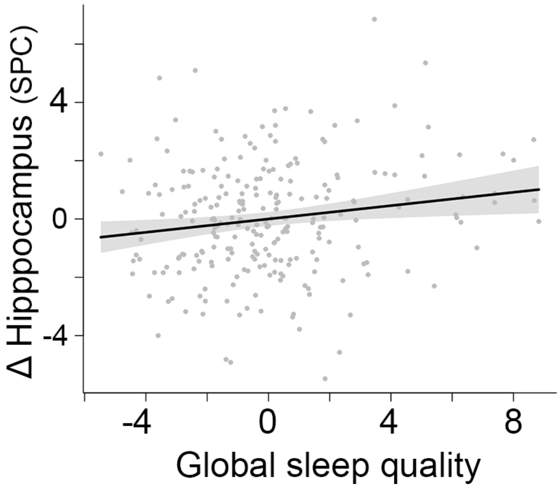

**Figure S3**. Self-reported sleep efficiency and polygenic scores
Self-reported sleep efficiency and polygenic scores (PGSs) for (A) sleep efficiency, and (B) AD.

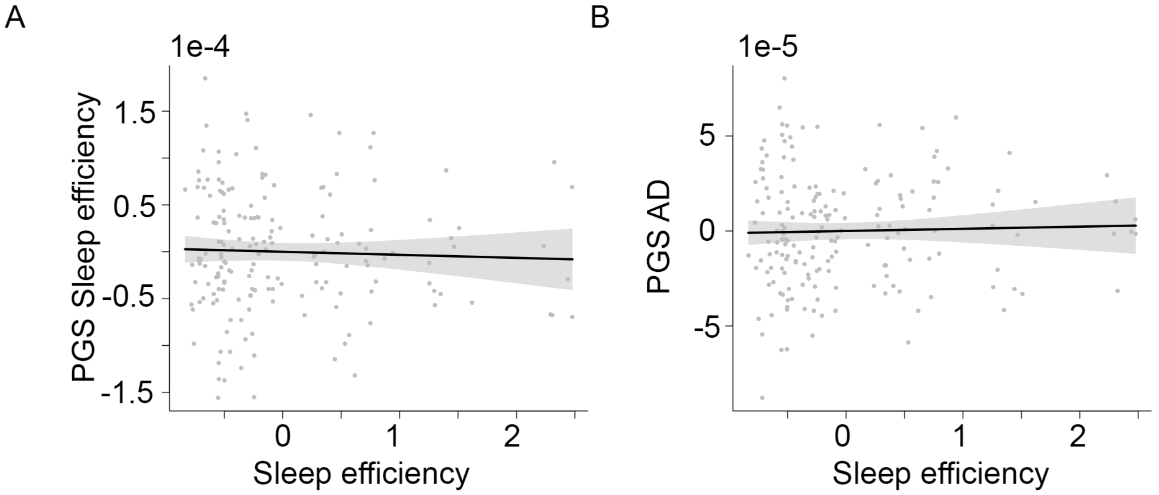

**Figure S4**. Hippocampal MD change and polygenic scores
Hippocampal MD change and polygenic scores (PGSs) for (A) sleep efficiency, and (B) AD.

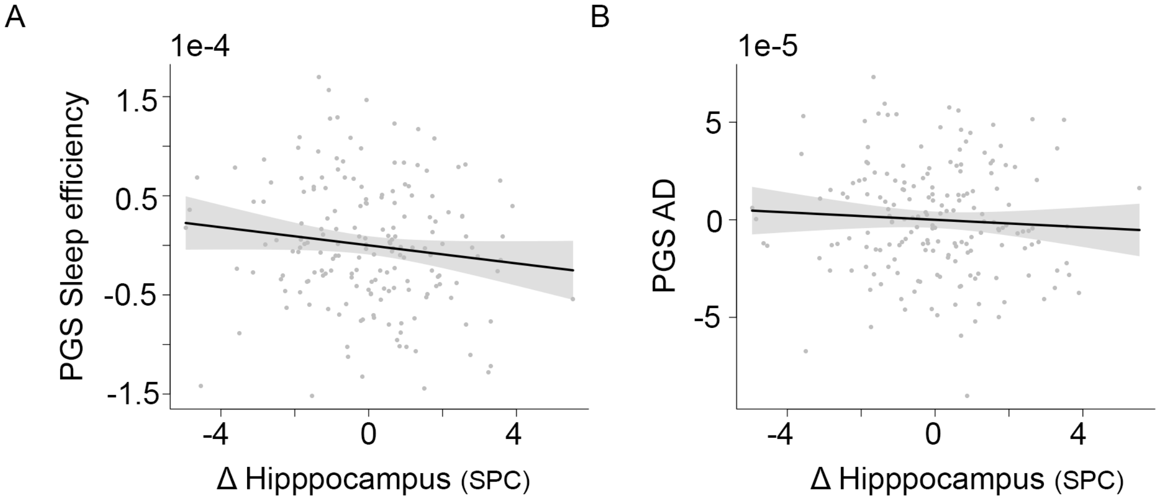

**Figure S5**. Distribution and classification of Aβ.

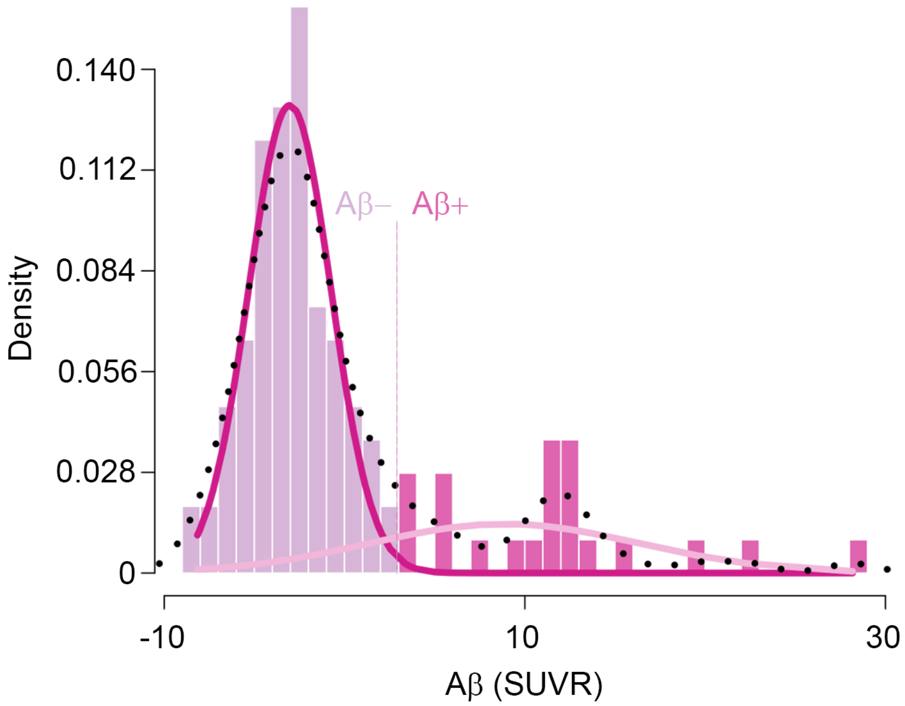

The best fit was a two-distribution solution (with unequal variance), represented in different colours. Fit of the distributions overlaid together with actual density (black dotted line). SUVR=standardized uptake value ratios. Aβ negative sample: n=85 (60%F), mean (SD, min-max) age: 67.4 (9.1, 44.4-80.8) years. Aβ positive sample: n=23 (48%F), mean (SD, min-max) age: 70 (6.6, 51.1-78.6) years.
